# Resting state functional connectivity in early post-anesthesia recovery is characterized by globally reduced anticorrelations

**DOI:** 10.1101/669457

**Authors:** Tommer Nir, Yael Jacob, Kuang-Han Huang, Arthur E. Schwartz, Jess W. Brallier, Helen Ahn, Prantik Kundu, Cheuk Y. Tang, Bradley N. Delman, Patrick J. McCormick, Mary Sano, Stacie G. Deiner, Mark G. Baxter, Joshua S. Mincer

## Abstract

Though a growing body of literature is addressing the possible longer-term cognitive effects of anesthetics, to date no study has delineated the normal trajectory of neural recovery due to anesthesia alone in older adults. We obtained resting state functional magnetic resonance imaging scans on 62 healthy human volunteers between ages forty and eighty before, during, and after sevoflurane (general) anesthesia, in the absence of surgery, as part of a larger study on cognitive function post-anesthesia. Resting state networks expression decreased consistently one hour after emergence from anesthesia. This corresponded to a global reduction in anticorrelated functional connectivity post-anesthesia, seen across individual regions-of-interest. Positively correlated functional connectivity remained constant across peri-anesthetic states. All measures returned to baseline 1 day later, with individual regions-of-interest essentially returning to their pre-anesthesia connectivity levels. These results define normal peri-anesthetic changes in resting state connectivity in healthy older adults.

## INTRODUCTION

Studying the effects of anesthetic agents on the brain in real time in the absence of surgery or a procedure is challenging since most anesthetics are performed to facilitate in conjunction with other procedures. However, the opportunity to image adults during anesthesia in the absence of surgery may enable deciphering the unique contribution of anesthetics to neural function. A key area of equipoise is how anesthesia impacts functional connectivity in the brain under general anesthesia. The full range of brain functional dynamics can be accessed through resting state functional magnetic resonance imaging (fMRI)^1–3^, thus enabling study of functional connectivity under anesthesia and comparison with awake states.^4–8^ This modality therefore offers the possibility of uncovering new patterns in resting state functional connectivity unique to anesthesia itself and discovery of imaging-based biomarkers predictive of individual or age-based variation which may be related to normal cognitive recovery and/or pathological states.

Such a mechanistic understanding of neural recovery from general anesthesia is vital to understanding the role, if any, of anesthetic agents in the pathogenesis of Perioperative Neurocognitive Disorders (PND)^9–13^. Around sixty-thousand patients undergo general anesthesia and surgery every day in the United States alone^14^, and it is currently unclear why some patients develop delirium, prolonged emergence, and delayed neurocognitive recovery lasting weeks to months. There is a preponderance of evidence suggesting that PND is not due to pharmacodynamics of the anesthetics themselves, but is more likely related to underlying medical conditions^15^ coupled with the persistent stress/inflammatory response elicited by surgery^16 17^. Ultimately, interpretation of studies focusing on PND require an understanding of the normal, expected trajectory of neural recovery in older adults, an essential yet missing piece.

The TORIE (Trajectory of Recovery in the Elderly) project^18^ is a prospective cohort study designed to delineate the normal trajectory of cognitive recovery after general anesthesia in the absence of surgery in healthy adults (40-80 years). A central part of the TORIE neuroimaging battery includes the acquisition of resting state functional magnetic resonance (rs-fMRI) scans. In the TORIE neuroimaging protocol, resting state scans are obtained at various time-points along the peri-anesthetic trajectory, including shortly prior to propofol induction, during a two-hour general anesthetic at a depth of one age-adjusted MAC (minimum alveolar concentration) of sevoflurane anesthesia, one-hour after emergence, and the following day. The resulting description of functional connectivity changes constituting a normal peri-anesthetic trajectory in healthy older adults will fill a gap in our current understanding and serve as a baseline for comparison with studies employing fMRI to investigate postoperative cognitive function in surgical patients.^19–22^ To this end, we report presently on analysis of rs-fMRI data for 62 healthy adult subjects.

## RESULTS

Subject demographics are summarized in Table 1. Independent component analysis (ICA)^23^ was employed to identify and characterize resting state network (RSN)^24^ activity. Region-of-interest (ROI)^25^) methods were employed to characterize changes in resting state functional connectivity along the peri-anesthetic trajectory.

**Table 1.** Subject demographics

|  | *n=62* | *standard deviation* |
| --- | --- | --- |
| *gender M/F* | 35/27 |  |
| *age (year)* | 58.87 | 11.49 |
| *MMSE (30 max)* | 28.74 | 0.98 |
| *education level (year)* | 15.32 | 2.07 |
| *BMI (kg/M^2)* | 25.72 | 3.09 |

### Resting state network expression

In order to characterize changes in RSN activity across the peri-anesthetic trajectory, group-ICA was applied to all 62 subjects at all 4 time-points along the peri-anesthetic trajectory (baseline (pre-anesthesia), anesthesia (ANES), post-anesthesia/early recovery (POST), and one day later (D1)). Multiple ICA components were aligned to known RSNs, specifically including somatosensory (SM), visual (VIS), default mode network (DMN), salience (SAL), dorsal attention (DA), frontoparietal (FP), language (LANG), cerebellar, and basal ganglia.

Specifically, RSN expression at each peri-anesthetic time-point was quantified as the volume of each RSN, measured as the number of voxels counted in the corresponding ICA component(s) (p<0.001 FDR corrected, cluster size p<0.05 FDR corrected)^26 27^. The dorsal attention RSN is highlighted in FIG 1. Changes in DA expression are illustrated qualitatively in FIG 1A, with the corresponding quantification by voxel count in FIG 1B. As compared to baseline, DA volume increased under ANES and decreased POST, returning to baseline levels at D1.

**Figure 1.**
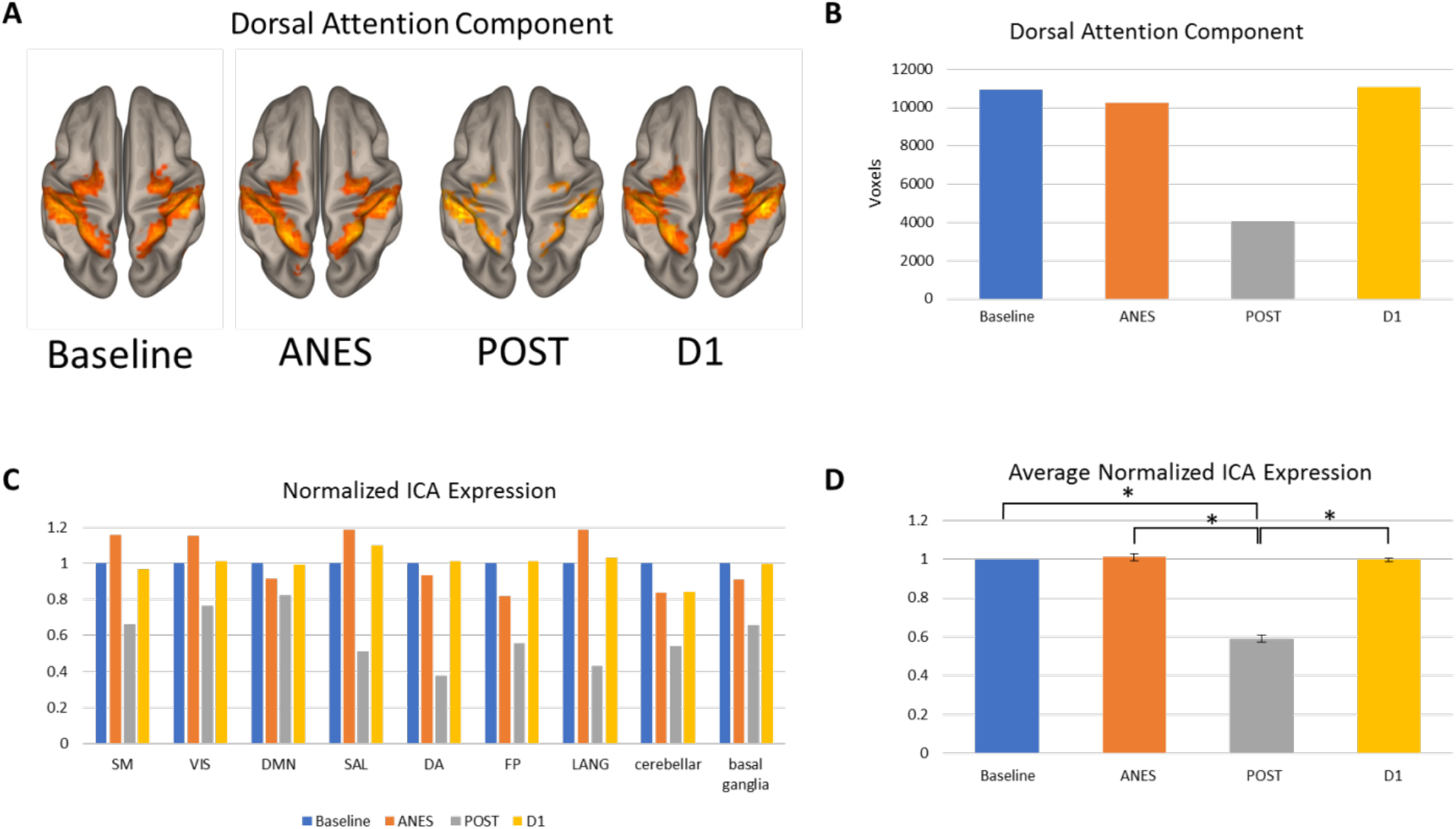
Resting state network expression. Volumetric analysis of ICA components matching canonical RSNs, before anesthesia (Baseline), during anesthesia (ANES), 1 hour after emergence (POST), and one day later (D1). (A) Dorsal attention network volume at the different time-points along the peri-anesthetic trajectory, illustrated (A) and quantified by voxel counts (B). Expression values for the various RSNs are shown in (C), with voxel counts at each time-point normalized to Baseline count value. The normalized values at each time-point are averaged across RSNs and shown in (D) (* denotes p-value<0.01, paired t-test).

Similar quantification of RSN volume for all RSNs studied is shown in FIG 2A. In order to facilitate comparison between networks, expression at each time point was normalized to each network’s baseline level. A consistent decrease in RSN expression was observed post-anesthesia. This is further captured by the mean normalized expression across networks (FIG 2B), illustrating an average 40% decrease POST (p<0.01). Average RSN expression under ANES did not change compared with baseline, though individual RSNs featured either an increase or decrease. For each RSN, expression levels returned to baseline at D1.

**Figure 2.**
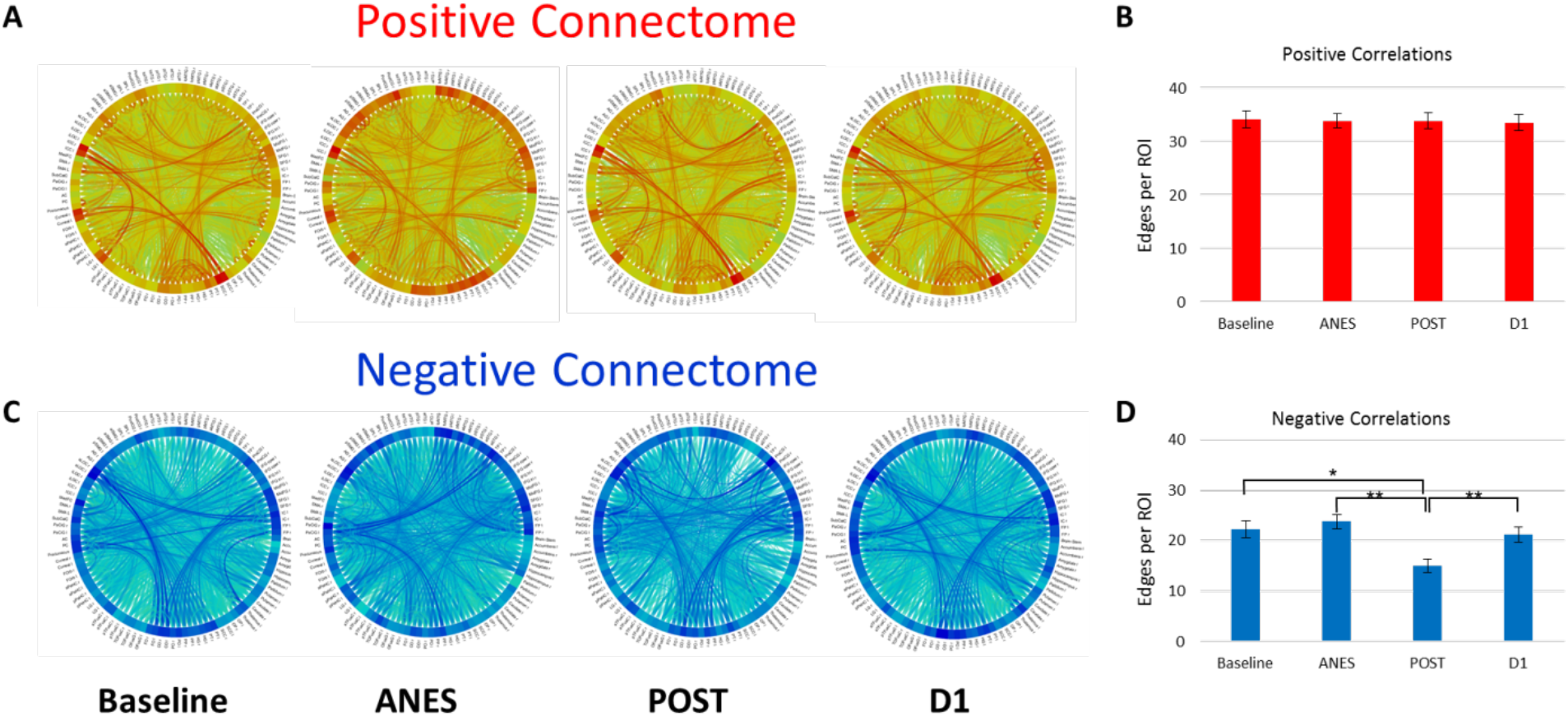
Correlated and anticorrelated functional connectivity. ROI-to-ROI analysis was performed using 106 cortical and subcortical ROIs (predefined in CONN). Functional connectivity was characterized as the number of statistically significant edges connecting ROIs and further divided into correlated and anticorrelated, represented by positive and negative edges respectively. Correlated (positive) edges are shown schematically for each time-point along the peri-anesthetic trajectory (A) and quantified as the average number of positive edges per ROI (B). Anticorrelated (negative) edges are similarly characterized in (C) and (D). Correlated functional connectivity does not change significantly across the peri-anesthetic trajectory. Changes in anticorrelated functional connectivity are significant: average anticorrelated edges per ROI increases significantly under anesthesia and decreases post-anesthesia as compared with Baseline (* denotes p<0.01 and ** denote p<10E-12, paired t-test).

### Functional connectivity

ROI-to-ROI analysis was performed using 106 predefined cortical and subcortical RSN masks (predefined within the CONN Functional Connectivity Toolbox^28^ and listed in Supplementary Table 1). Functional connectivity was quantified as the number of statistically significant edges connecting ROIs and divided into correlated (positive) and anticorrelated (negative) components and averaged as the number of edges (either correlated or anticorrelated) per ROI for a given time-point. Correlated (positive) functional connectivity was roughly the same across the peri-anesthetic trajectory. This is illustrated in FIG 2A, B. In contrast, anticorrelated connectivity changed significantly across the peri-anesthetic trajectory (FIG 2C, D). Specifically, averaged across all ROIs, the number of anticorrelated edges per ROI decreased by 31% in POST compared to baseline, (15 and 22 respectively, p=1.7 × 10^−12^) with return to baseline at D1. These changes in edge number are not due to signal dropout, as the average T-statistic of all ROIs is essentially unchanged across conditions (less than −4, see Supplementary Table 2).

Changes in individual ROIs were further explored by calculating the distribution of fractional change (versus baseline) at the other time-points. As seen in FIG 3, the observed average decrease in anticorrelations post-anesthesia was reflected at the single-ROI level, with nearly all ROIs demonstrating a negative fractional change at POST. There is greater variation under anesthesia, with some increasing and others decreasing (and some showing no change), with an overall slight positive bias. The return to baseline at day 1 observed for average connectivity values is reflected in the D1 distribution, which is centered rather tightly around zero.

**Figure 3.**
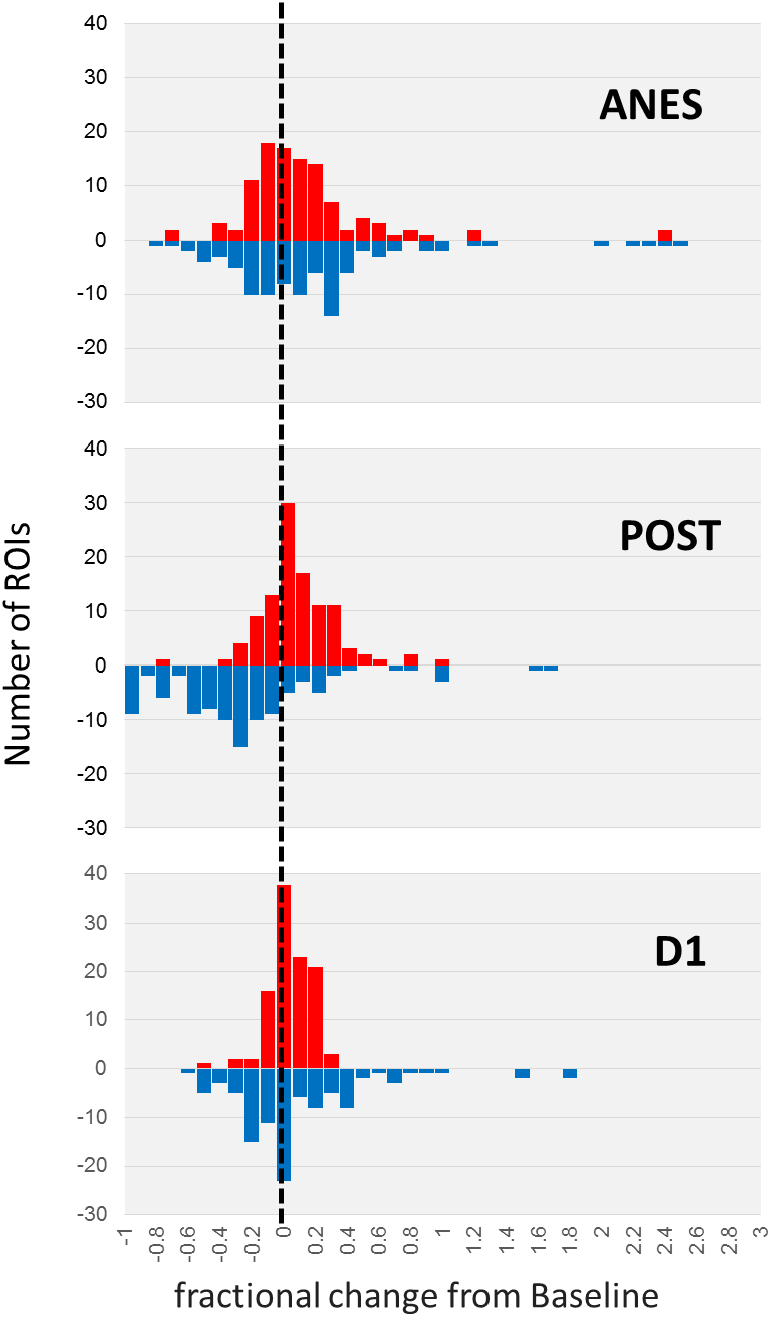
Distribution of individual ROI changes in correlated and anticorrelated functional connectivity. For each ROI, the fractional change in edge number from Baseline is calculated, and the resulting distribution (red for correlated and blue for anticorrelated connectivity) is plotted for (A) ANES versus Baseline, (B) POST versus Baseline, and (C) D1 versus Baseline.

The return to baseline at D1 is not an average phenomenon but rather represents each ROI essentially returning to its baseline connectivity value. FIG 4 demonstrates this explicitly, where the connectivity value at D1 is plotted against Baseline for each ROI (correlated connectivity in FIG 4A and anticorrelated connectivity in FIG 4B). The best-fit line for each curve is plotted as well, with intercept set at zero. For correlations the slope is 0.98 (R^2^ = 0.88) whereas for anticorrelations the slope is 0.91 (R^2^ = 0.71). The nearness of these slopes to 1 illustrates that the regression line is essentially the line D1 = Baseline.

**Figure 4.**
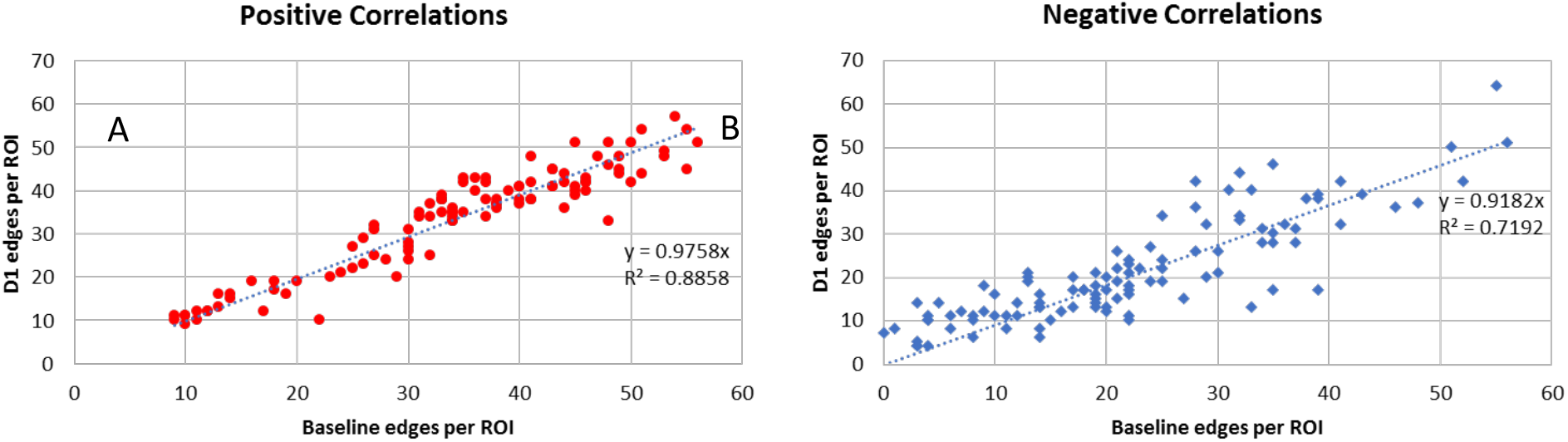
Return to Baseline connectivity at the single-ROI level. For each ROI, connectivity is plotted for D1 (y-axis) vs Baseline (x-axis) values (correlated connectivity in red (left) and anticorrelated connectivity in blue (right)). The best-fit regression line is plotted as well (with intercept set to zero). Slope is 0.98 (R^2^=0.89) for correlated connectivity and 0.92 (R^2^=0.72) for anticorrelated connectivity.

## DISCUSSION

Changes in RSN expression across the peri-anesthetic trajectory reflect global changes in anticorrelated functional connectivity. This is evident for connectivity measured in early post-anesthesia recovery (1 hour after emergence). At this time-point, RSN expression decreases across the board and a corresponding decrease in anticorrelations is seen in nearly all 106 anatomical ROIs. By contrast, changes at the individual ROI level under anesthesia are less uniform (some increasing, some decreasing) as compared to post-anesthesia. This variation is consistent with the variation in individual RSNs under anesthesia, with some increasing and others decreasing in expression (even as the mean change across RSNs is zero). Both measures of connectivity return to baseline one day later. This return to baseline is evident at the single-ROI level, with individual ROIs returning near their baseline connectivity values. Correlated functional connectivity remained constant on average across peri-anesthetic states.

Anticorrelations may help to differentiate and maintain boundaries between RSNs; classically this is represented by the interaction between the default mode network and task-positive networks.^29, 30^ Seen in this way, the global reduction in anticorrelated functional connectivity in early recovery is consistent with decreased RSN expression. The brain in early recovery/post-anesthesia features less-segregated, “fuzzy” networks. This characterization is consistent with the commonly observed clinical presentation of early recovery in which patients are conscious but clearly exhibit some transient cognitive deficits and suggests that globally-reduced anticorrelated connectivity may be seen as a functional neural correlate of early post-anesthesia recovery.

The reduced anticorrelations observed in early recovery may correspond to a similar signature seen in acute delirium states. A recent fMRI study of delirium subjects uncovered reduced anticorrelated functional connectivity between the posterior cingulate cortex (a component of the default mode network) and the dorsolateral prefrontal cortex (a component of the DA network) that was persistent even beyond clinical recovery.^31^ The authors of that study proposed that this finding may explain the cardinal symptoms of delirium such as impaired attention or consciousness. They further noted that this particular signature of reduced anticorrelated functional connectivity did not return to normal even after clinical recovery was noted. This is resonant with our findings: at our post-anesthesia time-point (one-hour post-emergence) the anesthetic agent is washed out, yet the signature of globally-reduced anticorrelations exists and does not return immediately to pre-anesthesia levels. The analogy between delirium and recovery from anesthesia is supported by a recent EEG study that uncovered similar functional connectivity signatures underlying both altered consciousness states.^32^ Because our participants did not exhibit delirium post-anesthesia emergence, it is possible that reduced anticorrelated connectivity is a permissive factor for the development of delirium.

Another recent study consistent with these findings demonstrated reduced anticorrelations between the default mode network and dorsal attention network in patients with mild cognitive impairment (MCI).^33^ Although the delirium and MCI studies focused specifically on default mode dorsal attention network interactions, we note that reduced anticorrelations observed in early recovery from general anesthesia are global and not limited to specific RSNs or ROIs. This suggests that previous studies that focused specifically on interactions of the default mode network with other RSNs identified a phenomenon that is expressed in these interactions but is not specific to them. It is possible that reduced anticorrelations are found more generally across other networks as well in delirium and MCI patients, a hypothesis that warrants further study (or reevaluation of existing data) in these populations.

In sum, the present work strongly supports the hypothesis that in healthy older adults, general anesthesia alone does not contribute to longer-term changes in neural activity. Observed measures of resting-state connectivity return to baseline within twenty-four hours of anesthesia exposure. Furthermore, the analogy between early recovery and delirium fostered by similar patterns in reduced anticorrelated functional connectivity has special significance. Beyond differentiating peri-anesthetic states, global anticorrelated functional connectivity conceivably may constitute an imaging-based biomarker that can predict whether an individual trajectory of recovery will veer into the pathology of post-operative delirium and cognitive dysfunction. Future research will explore these hypotheses.

## METHODS

### Participant selection

Results presented here were from first 62 healthy human volunteers who successfully completed the neuroimaging protocols of the TORIE project, representing just over 80% of anticipated total enrollment. This study was approved by the Institutional Review Board of the Icahn School of Medicine at Mount Sinai (New York, NY, USA, 212-824-8200) and registered at ClinicalTrials.gov (NCT02275026, Principal Investigators: Joshua Mincer, Mark Baxter, and Mary Sano, registered October 23, 2014).

Full details of participant enrollment, exclusion criteria, anesthesia and monitoring protocols, and experimental design may be found in the TORIE protocol paper.^18^ Briefly, all 62 volunteers were healthy (American Society of Anesthesiology physical class I or II) adults between the ages of 40 and 80 and had no underlying cognitive dysfunction as determined by baseline cognitive function testing (within the week prior to anesthesia exposure). Additionally, anatomical scans obtained prior to anesthesia induction were read by an on-site CAQ-credentialed neuroradiologist for evidence of intracranial pathology. Exclusion criteria were selected to ensure safety during anesthesia and MRI, the ability to complete testing at longer-term follow-up, and the absence of pathophysiology that could predispose to post-operative cognitive dysfunction, such as inflammatory conditions or cerebral microvascular disease. Exposure to general anesthesia within the last year was not a specific exclusion criterion. Demographics of the 62 subjects of the present study are found in Table 1.

### General anesthesia and monitoring protocols

The participants arrived early in the morning on the day they were to have general anesthesia. General anesthesia was induced following acquisition of the baseline TORIE neuroimaging battery, which included rs-fMRI and other modalities as well as the anatomical scans reviewed by the study neuroradiologist.

A baseline rs-fMRI scan was acquired prior to anesthesia induction. Following the awake scan, in preparation for induction of general anesthesia, the participant was positioned supine on the MRI gurney in the MRI suite, and a 22 gauge IV was placed. Standard ASA monitors were applied, and the participant was preoxygenated. Anesthesia was induced in the MRI suite with propofol 2 mg/kg IV, after which a laryngeal mask airway (LMA) was placed. Theoretically, if the LMA could not be seated properly, the procedure would have been aborted, though this was not an issue with any of the volunteers.

Anesthesia was maintained with inhaled sevoflurane at an age-adjusted depth of 1 minimum alveolar concentration (MAC). A bispectral index (BIS)^34^ level of 40-60 was obtained after LMA placement to aid in assessment of anesthetic depth during equilibration of inhaled sevoflurane and washout of propofol, after which the participant was returned to the MRI bore for scanning. End tidal sevoflurane concentration was used to measure anesthetic depth during scanning, along with physiological measures. Controlled ventilation was maintained to achieve a target ETCO2 of 30-35 mm Hg. Anesthesia was maintained for the next 2 hours, during which time three rs-fMRI scans (as well as other modalities employed in the TORIE neuroimaging protocol) were obtained. As necessary, the appropriate bolus administration of a pressor such as ephedrine (5 mg IV or 25 mg IM) or phenylephrine (100 µg IV) was occasionally administered by the anesthesiologist to maintain mean arterial blood pressure within 20% of baseline.

Following completion of the 2-hour anesthetic and scanning protocols, the participant was removed from the MRI bore and emerged from anesthesia. The LMA was removed when the participant awakened. Ondansetron (4 mg, IV) was given prior to emergence for antiemetic prophylaxis. No narcotics, benzodiazepines, steroids, or muscle relaxants were administered. The participant was then allowed to more fully awaken, all the time being monitored by the anesthesiologist.

The participant was returned to the MRI bore approximately 1 hour after anesthesia emergence, at which point additional rs-fMRI scanning was acquired. Note that by this point the participant exhibited normal and adequate spontaneous ventilation and as such would be normocapnic. Following this, the participant was transported to the post-anesthesia care unit (PACU) and further monitored until discharge. The participant returned the next morning (approximately 24 hours after anesthesia induction) for rs-fMRI acquisition.

### rs-fMRI acquisition and preprocessing

rs-fMRI scans were obtained at the following time-points along the peri-anesthetic trajectory: prior to propofol induction, during a two-hour anesthetic at a depth of 1 age-adjusted MAC (minimum alveolar concentration) of sevoflurane anesthesia, one-hour after emergence, and the following day (approximately 24 hours after anesthesia induction). For awake scans, participants were instructed to keep their eyes open and to let their thoughts wander.

rs-fMRI data were acquired with a multiband (MB) accelerated gradient multi-echo EPI sequence (FOV 224 × 224 mm, matrix 64 × 64, slice thickness 3.6 mm, 40 (32ch) slices for whole brain coverage, TR/TE = 1500/[10.8,28.68,46.56] ms, MB factor 2, blipped CAIPIRINHA phase encoding shift = FOV/3, bandwidth ∼ 1600 Hz/Pixel, echo spacing ∼0.5 ms, and total acquisition time ∼ 10 min (∼680 frames)).

Preprocessing of the rs-fMRI data employed the SAPIENT pipeline at the Icahn School of Medicine at Mount Sinai, which combines tools from the AFNI software suite^35^ in an automated fashion, including slice-timing correction, motion correction, and standard space normalization to an MNI template.^36–39^ The tools were applied in a manner appropriate for the processing of the multi-echo functional MRI data. For example, motion correction parameters were estimated from the first echo image and applied to images of the other echoes. Framewise displacement (FD) traces were computed for each fMRI dataset. Time points corresponding to FD>2mm were censored from all analyses. Degrees of freedom lost in censoring were accounted for in subsequent analysis. Additional nuisance regressors were generated as time courses computed from averaging fMRI signals of the white matter and CSF. These nuisance time courses were used as baseline regressors for subject-level analyses.

Subsequently, data analysis to determine components of blood-oxygen-level dependent (BOLD) signal was done using the meica.py Python software for the analysis of multi-echo fMRI data, implementing a pipeline involving preprocessing, high-dimensional independent components analysis, artifact component detection, and denoising by artifact removal. This software performs high-dimensional multi-echo independent components analysis (ME-ICA), and automatically separates BOLD and non-BOLD signals based on information on these respective signal types germane to multi-echo fMRI.^40–42^ Notably, the ME-ICA approach does not require the application of full-width-half-max spatial filters or temporal high pass filtering to attenuate noise in preprocessing, which is all handled in the main data analysis step involving separating BOLD from non-BOLD signals. ME-ICA denoising was performed on SAPIENT as well.

The ME-ICA denoised BOLD time-series for each rs-fMRI scan (i.e. each subject at each time-point along the peri-anesthetic trajectory) was further denoised using CompCor^43^ in order to fully remove any remaining non-BOLD respiratory artifact.^44^ Unlike other methods used for global signal removal (in particular Global Signal Regression (GSR)), CompCor does not introduce artefactual anticorrelations,^45^ a characteristic of particular importance when quantifying anticorrelated functional connectivity. CompCor denoising was carried out with the CONN Functional Connectivity Toolbox.

### rs-fMRI analysis

CONN^46^ was employed to analyze rs-fMRI data. CONN is an open-source Matlab/SPM-based cross-platform software that enables the computation, display, and analysis of functional connectivity fMRI data. All data presented is corrected for age. Independent Component Analysis (Group-ICA)^47^ was performed within CONN to obtain a set of ICA maps for all subjects at all time-points (before anesthesia (baseline), during anesthesia (ANES), 1 hour after emergence (POST), and one day later (D1)). An ICA map incorporates voxels with positively correlated BOLD time series. ICA components were mapped to known resting state networks (RSN) (salience, visual, dorsal attention, language, default mode network, somatosensory, and frontoparietal) which were subsequently visualized and further analyzed. In particular, RSN expression at each time-point was defined as the volume of the ICA component(s) matching each RSN. Volume was measured as the number of voxels counted in the corresponding ICA component (p<0.001 FDR corrected, cluster size p<0.05 FDR corrected). This allowed for comparison of each RSN expression across time-points, as well as mean expression for all RSNs.

Region of interest (ROI) analysis was also performed with CONN utilizing 106 cortical and subcortical ROIs predefined in CONN. ROI-to-ROI analysis calculated the functional connectivity between pairs of ROIs, defined as the Fisher-transformed bivariate correlation coefficients between each ROI’s BOLD time series. The ROI time series was computed by averaging voxel time series across all voxels within each ROI. Correlations above a determined significance threshold (p<0.01 FDR corrected) constituted edges between ROIs. These were further divided into correlated and anticorrelated edges, in which the sign of the correlation was either positive or negative respectively. Correlated and anticorrelated connectivity across all ROIs were compared at the 4 time-points.

Visualization of data in the figures employed images outputted from CONN itself and graphs created with Graphpad^48^. Microsoft Excel was also used for further statistical analysis of quantities outputted from CONN (for example calculation of various means and distributions).

## ACKNOWLEDGMENTS

We would like to thank our colleagues James Leader, Jacqueline Crittendon, Mohammed Ismail, Matthew Hartnett, Jessica Jong Kim, Carolyn Fan, Kirklyn Escondo, Rachelle Jacoby, Shanice Dumay, Johnny Ng, and Victoria Wang who all assisted immensely in the research. We gratefully acknowledge Dr Jeffrey Silverstein, who conceived of and implemented this study and served as principal investigator until his death. We acknowledge funding support via NIH/NIA R01AG046634. PJM, JWB, and JSM acknowledge support from NIH/NCI Cancer Center Support Grant P30 CA008748.

## TABLES

**Supplementary Table 1.** List of the 106 ROIs used in the ROI-to-ROI analysis. Coordinates in MNI152 space.

| **ROI** | **X** | **Y** | **Z** |
| --- | --- | --- | --- |
| Frontal Pole Right | 26.16 | 52.14 | 8.26 |
| Frontal Pole Left | -24.72 | 52.96 | 7.51 |
| Insular Cortex Right | 37.38 | 2.55 | -0.17 |
| Insular Cortex Left | -36.39 | 1.19 | 0.08 |
| Superior Frontal Gyrus Right | 14.67 | 18.43 | 56.96 |
| Superior Frontal Gyrus Left | -14.07 | 18.68 | 56.16 |
| Middle Frontal Gyrus Right | 39.12 | 18.62 | 42.79 |
| Middle Frontal Gyrus Left | -38.07 | 18.44 | 42.06 |
| Inferior Frontal Gyrus, pars triangularis Right | 51.87 | 27.76 | 7.71 |
| Inferior Frontal Gyrus, pars triangularis Left | -49.71 | 28.50 | 8.67 |
| Inferior Frontal Gyrus, pars opercularis Right | 52.21 | 15.42 | 16.20 |
| Inferior Frontal Gyrus, pars opercularis Left | -50.64 | 14.52 | 15.39 |
| Precentral Gyrus Right | 34.50 | -10.80 | 50.13 |
| Precentral Gyrus Left | -33.72 | -11.83 | 49.37 |
| Temporal Pole Right | 40.64 | 12.96 | -29.62 |
| Temporal Pole Left | -40.49 | 11.10 | -29.60 |
| Superior Temporal Gyrus, anterior division Right | 57.50 | -0.76 | -10.17 |
| Superior Temporal Gyrus, anterior division Left | -56.17 | -3.91 | -7.97 |
| Superior Temporal Gyrus, posterior division Right | 61.34 | -23.99 | 1.57 |
| Superior Temporal Gyrus, posterior division Left | -62.29 | -29.17 | 3.80 |
| Middle Temporal Gyrus, anterior division Right | 57.89 | -1.52 | -24.51 |
| Middle Temporal Gyrus, anterior division Left | -57.47 | -4.21 | -22.14 |
| Middle Temporal Gyrus, posterior division Right | 61.08 | -22.52 | -12.15 |
| Middle Temporal Gyrus, posterior division Left | -60.91 | -27.36 | -11.00 |
| Middle Temporal Gyrus, temporooccipital part Right | 58.18 | -49.22 | 1.60 |
| Middle Temporal Gyrus, temporooccipital part Left | -57.64 | -53.00 | 0.82 |
| Inferior Temporal Gyrus, anterior division Right | 46.23 | -2.41 | -41.11 |
| Inferior Temporal Gyrus, anterior division Left | -48.14 | -4.97 | -39.19 |
| Inferior Temporal Gyrus, posterior division Right | 53.42 | -23.46 | -28.13 |
| Inferior Temporal Gyrus, posterior division Left | -53.44 | -28.46 | -25.99 |
| Inferior Temporal Gyrus, temporooccipital part Right | 54.14 | -49.88 | -16.73 |
| Inferior Temporal Gyrus, temporooccipital part Left | -51.82 | -53.44 | -16.53 |
| Postcentral Gyrus Right | 37.62 | -26.37 | 52.64 |
| Postcentral Gyrus Left | -38.41 | -27.86 | 51.67 |
| Superior Parietal Lobule Right | 29.21 | -47.78 | 58.90 |
| Superior Parietal Lobule Left | -29.30 | -49.47 | 57.47 |
| Supramarginal Gyrus, anterior division Right | 58.42 | -27.06 | 37.81 |
| Supramarginal Gyrus, anterior division Left | -56.80 | -32.75 | 37.19 |
| Supramarginal Gyrus, posterior division Right | 55.21 | -40.37 | 33.60 |
| Supramarginal Gyrus, posterior division Left | -54.88 | -46.03 | 33.24 |
| Angular Gyrus Right | 51.93 | -51.80 | 32.36 |
| Angular Gyrus Left | -50.35 | -55.70 | 29.76 |
| Lateral Occipital Cortex, superior division Right | 32.97 | -71.12 | 38.92 |
| Lateral Occipital Cortex, superior division Left | -31.96 | -72.89 | 37.97 |
| Lateral Occipital Cortex, inferior division Right | 45.54 | -73.95 | -1.58 |
| Lateral Occipital Cortex, inferior division Left | -45.13 | -75.54 | -1.90 |
| Intracalcarine Cortex Right | 11.66 | -73.58 | 8.32 |
| Intracalcarine Cortex Left | -10.20 | -75.03 | 8.04 |
| Frontal Medial Cortex | 0.21 | 43.19 | -18.51 |
| Juxtapositional Lobule Cortex Right | 5.92 | -2.78 | 57.54 |
| Juxtapositional Lobule Cortex Left | -5.37 | -2.78 | 56.08 |
| Subcallosal Cortex | -0.07 | 20.54 | -14.83 |
| Paracingulate Gyrus Right | 6.55 | 36.57 | 22.69 |
| Paracingulate Gyrus Left | -6.21 | 36.65 | 20.79 |
| Cingulate Gyrus, anterior division | 0.80 | 18.29 | 24.35 |
| Cingulate Gyrus, posterior division | 0.78 | -36.62 | 29.98 |
| Precuneous Cortex | 0.96 | -59.29 | 38.03 |
| Cuneal Cortex Right | 8.85 | -78.55 | 27.89 |
| Cuneal Cortex Left | -8.22 | -80.29 | 27.14 |
| Frontal Orbital Cortex Right | 29.11 | 23.07 | -16.23 |
| Frontal Orbital Cortex Left | -29.54 | 23.66 | -16.57 |
| Parahippocampal Gyrus, anterior division Right | 22.36 | -8.05 | -30.26 |
| Parahippocampal Gyrus, anterior division Left | -21.87 | -9.11 | -30.30 |
| Parahippocampal Gyrus, posterior division Right | 22.90 | -30.53 | -16.76 |
| Parahippocampal Gyrus, posterior division Left | -21.90 | -32.43 | -16.89 |
| Lingual Gyrus Right | 13.57 | -63.49 | -4.96 |
| Lingual Gyrus Left | -12.27 | -65.67 | -5.44 |
| Temporal Fusiform Cortex, anterior division Right | 31.06 | -2.81 | -42.34 |
| Temporal Fusiform Cortex, anterior division Left | -31.88 | -4.43 | -41.90 |
| Temporal Fusiform Cortex, posterior division Right | 36.28 | -24.14 | -27.83 |
| Temporal Fusiform Cortex, posterior division Left | -35.97 | -29.53 | -25.08 |
| Temporal Occipital Fusiform Cortex Right | 35.05 | -50.06 | -16.64 |
| Temporal Occipital Fusiform Cortex Left | -33.50 | -53.68 | -15.97 |
| Occipital Fusiform Gyrus Right | 27.26 | -75.40 | -12.31 |
| Occipital Fusiform Gyrus Left | -26.58 | -76.57 | -13.59 |
| Frontal Operculum Cortex Right | 41.12 | 18.63 | 4.91 |
| Frontal Operculum Cortex Left | -39.70 | 18.33 | 4.53 |
| Central Opercular Cortex Right | 49.43 | -5.77 | 11.13 |
| Central Opercular Cortex Left | -47.99 | -8.63 | 11.81 |
| Parietal Operculum Cortex Right | 48.90 | -27.64 | 21.55 |
| Parietal Operculum Cortex Left | -48.36 | -31.85 | 20.46 |
| Planum Polare Right | 48.00 | -3.58 | -7.19 |
| Planum Polare Left | -46.61 | -5.97 | -7.34 |
| Heschl's Gyrus Right | 46.11 | -17.40 | 6.97 |
| Heschl's Gyrus Left | -45.20 | -20.32 | 7.19 |
| Planum Temporale Right | 54.97 | -25.07 | 12.07 |
| Planum Temporale Left | -52.70 | -29.71 | 10.79 |
| Supracalcarine Cortex Right | 8.22 | -74.49 | 14.08 |
| Supracalcarine Cortex Left | -8.36 | -73.25 | 14.80 |
| Occipital Pole Right | 17.73 | -95.13 | 8.31 |
| Occipital Pole Left | -16.85 | -96.50 | 6.74 |
| Thalamus right | 10.84 | -18.32 | 6.62 |
| Thalamus left | -9.99 | -19.22 | 6.29 |
| Caudate right | 13.30 | 10.01 | 10.49 |
| Caudate left | -12.79 | 8.98 | 9.74 |
| Putamen right | 25.50 | 1.78 | 0.30 |
| Putamen left | -24.90 | 0.48 | 0.34 |
| Pallidum right | 19.85 | -4.01 | -1.19 |
| Pallidum left | -18.96 | -5.12 | -1.33 |
| Hippocampus right | 26.50 | -20.96 | -14.25 |
| Hippocampus left | -25.18 | -23.19 | -13.81 |
| Amygdala right | 23.09 | -3.99 | -17.69 |
| Amygdala left | -23.00 | -4.95 | -17.73 |
| Accumbens right | 9.37 | 12.20 | -6.53 |
| Accumbens left | -9.46 | 11.50 | -7.17 |
| Brain-Stem | 0.48 | -29.73 | -34.98 |

**Supplementary Table 2.** Average T-statistic at each time-point for correlated and anticorrelated edges for the 106 ROIs.

|  | Baseline | ANES | POST | D1 |
| --- | --- | --- | --- | --- |
| Correlated | $7.24 \pm 0.46$ | $7.44 \pm 0.47$ | $6.5 \pm 0.39$ | $6.93 \pm 0.44$ |
| Anticorrelated | $-4.1 \pm 0.16$ | $-4.28 \pm 0.15$ | $-3.89 \pm 0.13$ | $-4.08 \pm 0.14$ |

## REFERENCES

1. Biswal B, Zerrin Yetkin F, Haughton VM, Hyde JS. Functional connectivity in the motor cortex of resting human brain using echo-planar MRI. Magn Reson Med 1995; 34: 537–41

2. Smith SM, Fox PT, Miller KL, et al. Correspondence of the brain’s functional architecture during activation and rest. Proc Natl Acad Sci 2009; 106: 13040–5

3. Lee MH, Smyser CD, Shimony JS. Resting-state fMRI: A review of methods and clinical applications. Am J Neuroradiol 2013; 34: 1866–72

4. Boveroux P, Vanhaudenhuyse A, Bruno MA, et al. Breakdown of within-and between-network Resting State Functional Magnetic Resonance Imaging Connectivity during Propofol-induced Loss of Consciousness. Anesthesiology 2010; 113: 1038–53

5. Liu X, Lauer KK, Ward BD, Li SJ, Hudetz AG. Differential effects of deep sedation with propofol on the specific and nonspecific thalamocortical systems: a functional magnetic resonance imaging study. Anesthesiology 2013; 118: 59–69

6. Akeju O, Loggia ML, Catana C, et al. Disruption of thalamic functional connectivity is a neural correlate of dexmedetomidine-induced unconsciousness. Elife 2014; 3: e04499

7. Bonhomme V, Vanhaudenhuyse A, Demertzi A, et al. Resting-state Network-specific Breakdown of Functional Connectivity during Ketamine Alteration of Consciousness in Volunteers. Anesthesiology 2016; 125: 873–88

8. Maier KL, McKinstry-Wu AR, Palanca BJA, et al. Protocol for the Reconstructing Consciousness and Cognition (ReCCognition) Study. Front Hum Neurosci 2017; 11: 284

9. Jevtovic-Todorovic V, Absalom AR, Blomgren K, et al. Anaesthetic neurotoxicity and neuroplasticity: an expert group report and statement based on the BJA Salzburg Seminar. Br J Anaesth 2013; 111: 143–51

10. Gleason LJ, Schmitt EM, Kosar CM, et al. Effect of Delirium and Other Major Complications on Outcomes After Elective Surgery in Older Adults. JAMA Surg 2015; 150: 1134–40

11. Fleisher LA. Brain Health Initiative: A New ASA Patient Safety Initiative. ASA Monit 2016; 80: 10–1

12. Inouye SK, Marcantonio ER, Kosar CM, et al. The short-term and long-term relationship between delirium and cognitive trajectory in older surgical patients. Alzheimer’s Dement 2016; 12: 766–75

13. Evered L, Eckenhoff R, Silbert B. Perioperative Cognitive Disorders Nomenclature and Comparative Data. Alzheimer’s Dement 2018; 14: 997

14. Brown EN, Lydic R, Schiff ND. General Anesthesia, Sleep, and Coma. N Engl J Med 2010; 363: 2638–50

15. Dokkedal U, Hansen TG, Rasmussen LS, Mengel-From J, Christensen K. Cognitive Functioning after Surgery in Middle-aged and Elderly Danish Twins. Anesthesiology 2016; 124: 312–21

16. Vacas S, Degos V, Feng X, Maze M. The neuroinflammatory response of postoperative cognitive decline. Br Med Bull 2013; 106: 161–78

17. Eckenhoff RG, Laudansky KF. Anesthesia, surgery, illness and Alzheimer’s disease. Prog Neuropsychopharmacol Biol Psychiatry Univ Penn, Dept Anesthesiol & Crit Care, Perelman Sch Med, Philadelphia, PA 19104 USA; 2013; 47: 162–6

18. Mincer JS, Baxter MG, Brallier JW, Sano M, Schwartz AE, Deiner SG. Delineating the Trajectory of Cognitive Recovery From General Anesthesia in Older Adults: Design and Rationale of the TORIE (Trajectory of Recovery in the Elderly) Project. Anesth Analg 2018; 126: 1675–83

19. Abu-Omar Y, Cader S, Guerrieri Wolf L, Pigott D, Matthews PM, Taggart DP. Short-term changes in cerebral activity in on-pump and off-pump cardiac surgery defined by functional magnetic resonance imaging and their relationship to microembolization. J Thorac Cardiovasc Surg 2006; 132: 1119–25

20. McDonagh DL, Berger M, Mathew JP, Graffagnino C, Milano CA, Newman MF. Neurological complications of cardiac surgery. Lancet Neurol 2014; 13: 490–502

21. Price CC, Tanner JJ, Schmalfuss I, et al. A pilot study evaluating presurgery neuroanatomical biomarkers for postoperative cognitive decline after total knee arthroplasty in older adults. Anesthesiology 2014; 120: 601–13

22. Browndyke JN, Berger M, Harshbarger TB, et al. Resting-State Functional Connectivity and Cognition After Major Cardiac Surgery in Older Adults without Preoperative Cognitive Impairment: Preliminary Findings. J Am Geriatr Soc 2017; 65: e6–12

23. Calhoun VD, Adali T, Pearlson GD, Pekar JJ. A Method for Making Group Inferences from Functional MRI Data Using Independent Component Analysis. 2001; 151: 140–51

24. Damoiseaux JS, Rombouts SA, Barkhof F, et al. Consistent resting-state networks across healthy subjects. Proc Natl Acad Sci U S A 2006; 103: 13848–53

25. Poldrack RA. Region of interest analysis for fMRI. Soc Cogn Affect Neurosci 2007; 2: 67–70

26. Case M, Zhang H, Mundahl J, et al. Characterization of functional brain activity and connectivity using EEG and fMRI in patients with sickle cell disease. NeuroImage Clin; 2017; 14: 1–17

27. Noirhomme Q, Soddu A, Lehembre R, et al. Brain Connectivity in Pathological and Pharmacological Coma. Front Syst Neurosci 2010; 4: 1–6

28. Whitfield-Gabrieli S, Nieto-Castanon A. Conn: A Functional Connectivity Toolbox for Correlated and Anticorrelated Brain Networks. Brain Connect 2012; 2: 125–41

29. Greicius M, Krasnow B, Reiss A, Menon V. Functional connectivity in the resting brain: A network analysis of the default mode hypothesis. Proc Natl Acad Sci U S A 2003; 100: 253–8

30. Fox MD, Snyder AZ, Vincent JL, Corbetta M, Essen DC Van, Raichle ME. The human brain is intrinsically organized into dynamic, anticorrelated functional networks. Sci York 2007; 102: 9673–8

31. Choi SH, Lee H, Chung TS, et al. Neural network functional connectivity during and after an episode of delirium. Am J Psychiatry 2012; 169: 498–507

32. Numan T, Slooter AJC, Kooi AW Van Der, et al. Functional connectivity and network analysis during hypoactive delirium and recovery from anesthesia. Clin Neurophysiol 2017; 128: 914–24

33. Esposito R, Cieri F, Chiacchiaretta P, et al. Modifications in resting state functional anticorrelation between default mode network and dorsal attention network?: comparison among young adults, healthy elders and mild cognitive impairment patients. Brain Imaging and Behavior 2018; 12 127–41

34. Sigl JC, Chamoun NG. An introduction to bispectral analysis for the electroencephalogram. J Clin Monit 1994; 10: 392–404

35. Cox RW HJS. AFNI: Software for analysis and visualization of functional magnetic resonance neuroimages. Comput Biomed Res 1996; 29: 162–73

36. Avants B, Tustison N, Song G. Advanced Normalization Tools (ANTS). Insight J 2009; 1–35

37. Jenkinson M, Beckmann CF, Behrens TEJ, Woolrich MW, Smith SM. Fsl. Neuroimage 2012; 62: 782–90

38. Cox RW HJS. AFNI: Software for analysis and visualization of functional magnetic resonance neuroimages.. Comput Biomed Res 1996; 29: 162–73

39. Fischl B, Dale AM. Measuring the thickness of the human cerebral cortex from magnetic resonance images. Proc Natl Acad Sci 2000;

40. Kundu P, Inati SJ, Evans JW, Luh WM, Bandettini PA. Differentiating BOLD and non-BOLD signals in fMRI time series using multi-echo EPI. Neuroimage 2012; 60: 1759–70

41. Kundu P, Brenowitz ND, Voon V, et al. Integrated strategy for improving functional connectivity mapping using multiecho fMRI. Proc Natl Acad Sci U S A 2013; 110: 16187–92

42. Kundu P, Voon V, Balchandani P, Lombardo M V, Poser BA, Bandettini P. Multi-Echo fMRI: A Review of Applications in fMRI Denoising and Analysis of BOLD Signals. Neuroimage 2017; 154: 59–80

43. Behzadi Y, Restom K, Liau J, Liu TT. A component based noise correction method (CompCor) for BOLD and perfusion based fMRI. Neuroimage 2007; 37: 90–101

44. Power JD, Plitt M, Gotts SJ, et al. Ridding fMRI data of motion-related influences: Removal of signals with distinct spatial and physical bases in multiecho data. Proc Natl Acad Sci 2018; 115: 201720985

45. Chai XJ, Castañán AN, Öngür D, Whitfield-Gabrieli S. Anticorrelations in resting state networks without global signal regression. Neuroimage 2012;

46. Whitfield-Gabrieli S, Nieto-Castanon A. *Conn*: A Functional Connectivity Toolbox for Correlated and Anticorrelated Brain Networks. Brain Connect 2012;

47. Calhoun VD, Adali T, Pearlson GD, Pekar JJ. A Method for Making Group Inferences from Functional MRI Data Using Independent Component Analysis V.D. Hum Brain Mapp 2001; 140–51

48. GraphPad Software Inc. Graphpad Prism. GraphPad Prism. Data Anal. Sci. Graph. 2010.

